# Double-stranded RNA induces retinal pigment epithelium cell degeneration and inflammation

**DOI:** 10.1101/2024.03.11.584455

**Authors:** Garrett Klokman, YongYao Xu, Kyle Bond, Xiaoqiu Wu, Joshua Schustak, Jorgi Mandelbaum, Michael Twarog, Hongwei Han, Mary-Kate Paulina, Matthew Coble, Christopher Hayden, Jean-Rene Galarneau, John Demirs, Yubin Qiu, Robert Esterberg, Qian Huang, Ganesh Prasanna, Magali Saint-Geniez, Jorge Aranda, Yi Bao

## Abstract

RIG-I signaling has been previously implicated as a driver of inflammation to the retinal pigment epithelium (RPE) during age-related macular degeneration (AMD). Double-stranded RNA (dsRNA) is known to initiate RIG-I signaling and lead to a type I interferon response. We show through shRNA knockdown that RIG-I is essential for initiating an interferon response in iPS-RPE in response to both synthetic dsRNA-mimetic 3p-hpRNA and the double-stranded retrotransposable element *Alu*. Analysis of human tissue from patients suffering from AMD show accumulation of dsRNA, peaking at the geographic atrophy (GA) stage. Using a new murine model of 3p-hpRNA subretinal challenge to RPE cells, we confirmed that accumulation of dsRNA initiates a type I interferon response, as well as RPE and photoreceptor degeneration. Although RPE response to synthetic dsRNA was acute, extensive leukocyte migration was observed. The results from this study verify the importance of RIG-I signaling in regulating inflammation in the subretinal space and implicates dsRNA accumulation as a driver of AMD pathogenesis.

## Introduction

Age-related macular degeneration (AMD), a disease that leads to progressive visual impairment and blindness, is characterized by pro-oxidative and pro-inflammatory stress to the retinal pigment epithelial cells (RPE) culminating in the death of both RPE and photoreceptors [1]. Despite recent advancements in the treatment of late-stage dry AMD, preventing intermediate AMD progression remains an unmet therapeutic need. Several mechanistic and clinical studies of AMD pathology have identified an important role of the immune system in disease progression driven by RPE cell stress, including both complement and type I interferon [2]. Targeting complement activation has led to the first approved treatment for geographic atrophy (GA), the final stage of dry AMD [3]. The role of type I interferon has been shown to be important for AMD progression as well, but finding a safe and efficacious target within the pathway has remained challenging in absence of a disease-replicating animal model [4, 5].

Our group has identified retinoic acid-inducible gene I (RIG-I) as the key driver of the type I interferon response in RPE [6]. RIG-I is an intracellular receptor that acts as a sensor for double-stranded RNA (dsRNA). Known as the “antiviral response”, dsRNA binding to RIG-I initiates a signaling cascade leading to NF-κB-mediated expression and secretion of type I interferons such as IFNβ. In addition to our findings, others have reported the accumulation of retrotransposons in both AMD and other aging disorders [7, 8]. *Alu* elements, the most abundant transposable elements found throughout the human genome, mediate gene transcription, gene splicing, and protein translation [9]. Interestingly, the *Alu* elements inherently form double-stands due to their repetitive sequence makeup [10]. These findings indicate that dsRNA may play an integral role in driving the pro-inflammatory component of AMD pathogenesis.

The goal of this work is to better understand the importance of RIG-I and dsRNA in AMD. We investigated the presence of dsRNA in RPE during AMD from human tissue samples, and explored whether RIG-I is essential for mediating an inflammatory response to dsRNA *in vitro* using iPS-RPE. Importantly, we also report that delivery of dsRNA to RPE *in vivo* using a novel model could induce an acute interferon response, resulting in RPE degeneration and leukocyte recruitment into the retina. These findings reinforce the importance of RIG-I signaling in RPE stress and confirms the detrimental effects of dsRNA in AMD.

## Material and Methods

### Cell Culture

iPS-derived RPE cell line (iPS-RPE) was purchased from FUJIFILM Cellular Dynamics (Madison, WI). Cells were cultured in Lonza RtEGM medium (Lonza) at 37°C and 5% CO_2_. iPS-RPE cells were cultured on tissue culture treated plates, or on 24-well 0.4 μm transwells (Corning, #353095) transmembrane inserts for polarization (Corning, Corning, NY; 353095). Cells were cultured at least 3 weeks post seeding to form mature monolayers. Maturation of cells was confirmed by pigmentation, immunofluorescence staining for ZO-1 (Thermo Fisher Scientific, Waltham, MA #33-9111) and BEST1 (Novus, #NB300-164), and transepithelial resistance (TER).

### Gene Knockdown

Lentivirus encoding shRNA were made by VectorBuilder (Chicago, IL). Viral transduction of matured iPS-RPE were aided using TransDux virus transduction reagent (System Biosciences, cat# LV850A-1) and spin infection (800 g, 1 h at 32ºC). Knockdown efficacy was validated at the mRNA or protein level by either RT-qPCR or Western blot one week post-transduction.

### Nucleic acid labeling

In indicated experiments, 3p-hpRNA (InvivoGen, tlrl-hprna) was fluorescently labeled using Ulyssis Alexa Fluor 647 Nucleic Acid Labeling Kit (Thermo Fisher, U21660). In short, lyophilized 3p-hpRNA was reconstituted in Labeling Buffer, then labeled with Alexa Fluor 647 following manufacturer guidelines. Labeled 3p-hpRNA (AF647-3p-hpRNA) was purified using Zymo Direct-zol Miniprep Kit (Zymo Research, R2051) and eluted in physiologic water (InvivoGen, tlrl-hprna). Preparations were confirmed labeled through visual inspection of solution and through Nanodrop quantification (ThermoFisher).

### dsRNA Transfection

Both positive and negative strands of *Alu* RNA were produced through *in vitro* transcription from corresponding plasmids using T7 RNA polymerase and annealed. Annealed dsRNA was used for iPS-RPE transfection with Lipofectamine™ 3000 Transfection Reagent (L3000001) according to manufacturer instructions up to a final concentration of 40 ng/ml. Annealing of *Alu* RNA into double-stranded RNA (*dsAlu*) was confirmed via immunostaining of cells using a dsRNA specific antibody (J2, Scions) after transfection.

For AF647-3p-hpRNA solutions, confirmation of transfection potential was performed on iPS-RPE cells using NeuroPORTER (Sigma #NPT01). In short, AF647-3p-hpRNA was complexed with NeuroPORTER following manufacturers guidelines. Complexes were added to 24-well transwells of mature iPS-RPE up to a final concentration of 100 ng/ml and incubated for 24 h before imaging on EVOS M7000 (Thermo-Fisher).

### Transepithelial resistance (TER) measurement

Human RPE cells were seeded (200 k cells/cm^2^) in 24-well transwells (Corning #353,095) and allowed to expand and mature for at least 3 weeks in serum-free RtEBM media. After reaching maturation, barrier function was assessed by monitoring TER every 30 min for 18 h by means of a cellZscope 2 (NanoAnalytics GmbH, Münster, Germany). The resistance values for individual well at specific times (Ω/cm2) were determined, subtracted for background resistance produced by the blank filter and culture medium (as 0%), and normalized to baseline resistance prior to stimulation (as 100%).

### RT-qPCR

Messenger RNAs were isolated from iPS-RPE using TurboCapture 96 mRNA Kit (Qiagen). For mouse tissue, RNA was isolated using RNeasy Plus Mini Kit (Qiagen #74,136). RNA was reverse transcribed using the High-Capacity cDNA Reverse Transcription Kit (Applied Biosystems, Foster City, CA). Real-time PCR was performed using FAM-labeled TaqMan probes targeting genes of interest and a VIC-labeled TaqMan probe targeting *β-actin, GAPDH* (for *in vitro*) or *18s* (for *in vivo*) as control (Applied Biosystems). Reactions were run with TaqMan Universal PCR Master Mix on the ViiA7 system (Applied Biosystems) according to the instructions of the manufacturer.

### Human donor sample collection

Postmortem human eyes were procured from the Lions World Vision Institute (LWVI; Tampa FL) with consent of donors or donors’ next of kin and in accordance with the Eye Bank Association of America medical standards, US/Florida law for human tissue donation, the Declaration of Helsinki and Food and Drug Administration (FDA) regulations, and Novartis human tissue registration working practice guidelines regarding research using human tissues. All study tissues were preserved within 6 h postmortem or less. AMD grading on hematoxylin and eosin (H&E) sections was determined following criteria from the Sarks, Age-Related Eye Disease Study (AREDS), and The Beckman Group clinical grading systems [11-13]. We classified those donors into 3 groups, including patients with GA (AMD4) and age-matched controls (AMD1), and intermediate AMD (iAMD, combined AMD2 and AMD3). The donor information used for this study is listed in Supplemental Table 1.

### Immunohistochemistry

Eyes were injected intravitreally with 100 μl of Modified Davidson’s Fixative (MDF, H0290-500ML, Sigma-Aldrich, St. Louis, MO) and preserved as whole globes in that same fixative for 48 h, followed with 70% alcohol for an additional 48 h at room temperature. The eyes were embedded in paraffin wax, serially sectioned at a thickness of 5 μm through the optic nerve, and processed for dsRNA antibody staining (J2, Scions) using Leica Bond RX according to manufacturer’s instructions. 20X and 40X images were taken by the Aperio AT2 scanner. IgG antibody was used as a negative control. In each stained section (one per J2 antibody per eye), immunoreactivities were scored semiquantitatively as 0 (negative), 1 (weak), 2 (moderate), 3 (strong), or 4 (very strong).

### Gene Array

Custom TaqMan Array Cards were ordered from ThermoFisher (cat# 4391526). Total RNA (1 μg/sample) was reverse transcribed to cDNA using SuperScript™ IV VILO™ Master Mix (ThermoFisher, cat# 11756050). A fraction of cDNA was mixed with TaqMan qPCR Master Mix (ThermoFisher, cat# 4444557), loaded into fill reservoir of the card and the reaction run on the ViiA™ 7 instrument. The results were analyzed using relative quantification (ΔΔCt) method, normalized to internal controls.

### 3p-hpRNA subretinal injection

All procedures and housing conditions were approved by Novartis Cambridge Institutional Animal Care and Use Committee. Control and dsRNA-treated eight-weeks-old female C57/Bl6J mice (Jackson) were injected subretinally (SR) with 1 μL of the transfection reagent Neuroporter (Millipore, NPT01) alone or in combination with 3p-hpRNA (Invivogen, Cat tlrl-hpRNA), respectively. Both injection preparations contained 1:50 sodium fluorescein (Alcon, Cat # 00065009265). Pupils were dilated with topical application of 1% cyclopentolate and 2.5%–10% phenylephrine. Mice were subsequently anesthetized with an intraperitoneal injection of ketamine/xylazine cocktail (100/10 mg/kg) and 0.5% proparacaine was applied to the eyes. Eye moisture was maintained by application of Genteal (Novartis). An initial puncture was made nasally through the sclera, posterior to the limbus, with a 30-gauge-needle. SR injection was performed using a blunt-ended 33-gauge needle fitted on a 10 μL Hamilton syringe and inserted tangentially under the lens through the scleral incision. Successful SR injections were confirmed by visualization of a fluorescein-containing bleb at the time of injection. Tobrex (Alcon, Fort Worth, TX) was applied to the eyes following injection. Mice were monitored for recovery for 24 h and maintained until takedown.

### Posterior eye cup immunostaining, imaging and analysis

Eyes were dissected from control and mice challenged with dsRNA. Next, the cornea, lens and retina were excised leaving the posterior eye cup (PEC) consisting of the RPE, choroid and sclera. The PEC was then bleached by incubating in 10% hydrogen peroxide for 2 h at 65ºC. The PEC was washed thrice in PBS (5 min each) at room temperature (RT). Non-specific binding was inhibited by incubating for 2 h in blocking buffer (5% BSA or Goat Serum (Invitrogen, #500622) + Triton X-100 (VWR, 97062-208)). Mouse anti-ZO-1 Alexa Fluor 647 conjugated antibody (1:100, ThermoFisher MA3-39100-A647) diluted in blocking buffer containing Hoechst 33342 (1:1000, ThermoFisher 33342) was added to the tissue for 48 h at 4ºC with gentle rocking. For IBA1 staining, PECs were then stained overnight at 4ºC with IBA1 antibody (1:1000, Wako 019-19741) in blocking buffer followed by PBS wash 15 min three times and then incubation for 2 h at room temperature with Alexa Fluor 594 conjugated donkey anti-rabbit antibody (1:1000, ThermoFisher A-21207). Stained samples were then washed for 15 min three times in PBS. PECs were then opened with four incisions forming petal-shaped lobes. The PEC was flatmounted with glass coverslips and mounting medium (Invitrogen, TA-030-FM). The stained samples were imaged on a ZEISS LSM 900 with Airyscan. Samples were tile imaged at 10X to capture the entire flatmount (for IBA1) or 2.5X to find the initial incision point opposite the optic nerve (for ZO-1). Identified injection sites were further imaged at 10X and 20X in a 982.8 μm x 982.8 μm square and excluded from analysis to avoid injection related artifacts. 5 images per PEC were taken directly outside of the excluded area at 40X. For RPE morphological assessment, three double-blinded graders scored the RPE morphology of several regions across 10 animals per group based on deviation from reference images. For RPE morphometric analysis, a novel Arivis software pipeline specifically designed to assess RPE ZO-1 staining was used to determine cell roundness.

### Transmission electron microscopy

Female C57BL/6 mice were injected SR with 1 uL vehicle transfection agent (controls) or with 3p-hpRNA dsRNA. Two control animals and two 3p-hpRNA dosed animals were selected for transmission electron microscopy (TEM) investigation. For each animal, the eye was enucleated with the superior aspect of the eye marked with histochemical dye to establish orientation, and fixed by immersion in Modified Karnovsky’s Fixative (2% Paraformaldehyde and 2.5% Glutaraldehyde in 0.1 M Sodium Cacodylate Buffer, pH 7.4 (Electron Microscopy Sciences (Hatfield, PA), #15960-01). The cornea, lens, and iris were removed. Using a stereoscope, the subretinal injection site was visualized and the eye was bisected adjacent to it. Section containing the injection site was post-fixed in 1% osmium tetroxide in 0.1 M cacodylate buffer, en-block stained with aqueous 1% uranyl acetate, and dehydrated in an upgraded acetone series using Pelco BioWave tissue processor (Ted Pella, INC. Redding, CA). The samples were embedded in Quetol-modified Spurr’s epoxy at 60°C for 48 h. Samples were rough trimmed until the first complete section of retina was visible. Brightfield evaluation of ∼0.5μm thick toluidine-blue stained sections was performed every ∼50μm until reaching the target area. Ultrathin sections (∼70 nm thick) were generated on a Leica Ultracut UCT ultramicrotome, collected on 300 mesh formvar-coated grids and double stained with Reynolds lead citrate and 1% aqueous uranyl acetate. TEM was performed on an FEI/Thermo Tecnai T12 Spirit BioTwin TEM with an Olympus/SIS digital camera system. Retinal regions distant from the injection site (approximately 300 to 500 μm away from the subretinal injection site) were used as reference controls to avoid areas affected by any procedure-related effects.

### Data Analysis

All *in vitro* RT-qPCR experiments and TER results were statistically analyzed using Student’s t-test. All *in vivo* RT-qPCR experiments were statistically analyzed using Dunnett’s one-way ANOVA. For immunostaining analyses, ranked data was analyzed with Mann-Whitney U test, while ARIVIS quantification raw data was compared using Pearson’s correlation analysis. All statistics and graphs were generated using Graphpad Prism 9.4.1 (Dotmatics).

## Results

### ISG15 expression in AMD is dependent on RIG-I

Exposure to intracellular dsRNA led to increased expression of IFNβ, indicating a type I interferon response in ARPE-19 and iPS-RPE, and increased susceptibility to a degenerative phenotype [6]. IFNβ secretion promotes lymphocyte recruitment and activation, which can lead to the secretion of inflammatory and pro-degenerative cytokines such as TNFα [14]. Additionally, autocrine IFNβ signaling can initiate an autoregulatory feedback loop, as indicated by increased ISG15 expression [6].

Previously we have validated RIG-I as essential to induce an inflammatory response, such as secretion of IFNβ and ISG15 expression, to dsRNA stress using CRISPR/Cas9 mediated knockdown of RIG-I in ARPE19 cells [6]. Because ARPE-19 have limitations in their ability to replicate all RPE characteristics, such as the inability to form proper tight junctions, we investigated if these findings were replicable in iPS-RPE cells. The synthetic dsRNA-mimetic 3p-hpRNA, which contains a dsRNA-like hairpin-loop structure detectable by dsRNA sensors [15], increases *RIG-I* expression in iPS-RPE after transfection (Figure 1A-B). Using shRNAs against *RIG-I*, we were able to stably repress *RIG-I* expression for up to 4 weeks (shRNA1 = 78.6% reduction, shRNA2 = 97.7% reduction) (Figure 1A). Replicating the results from our studies with ARPE-19, dsRNA could profoundly induce *ISG15* expression (Figure 1B). Knockdown of *RIG-I* successfully repressed *ISG15* induction by dsRNA, validating our proposed signaling paradigm (Figure 1B). Interestingly, induction of RIG-I signaling via 3p-hpRNA significantly decreased the expression of the RPE-specific visual cycle protein RPE65 (Fig. 1C), suggesting cellular de-differentiation and loss-of-function. Epithelial barrier function, a major phenotypic characteristic of RPE health and functionality, was measured through trans-epithelial resistance (TER) in response to dsRNA stimulation. While dsRNA induces a reduction in barrier function, knockdown of RIG-I was able to rescue cells (Figure 1D-E).

**Figure 1:**
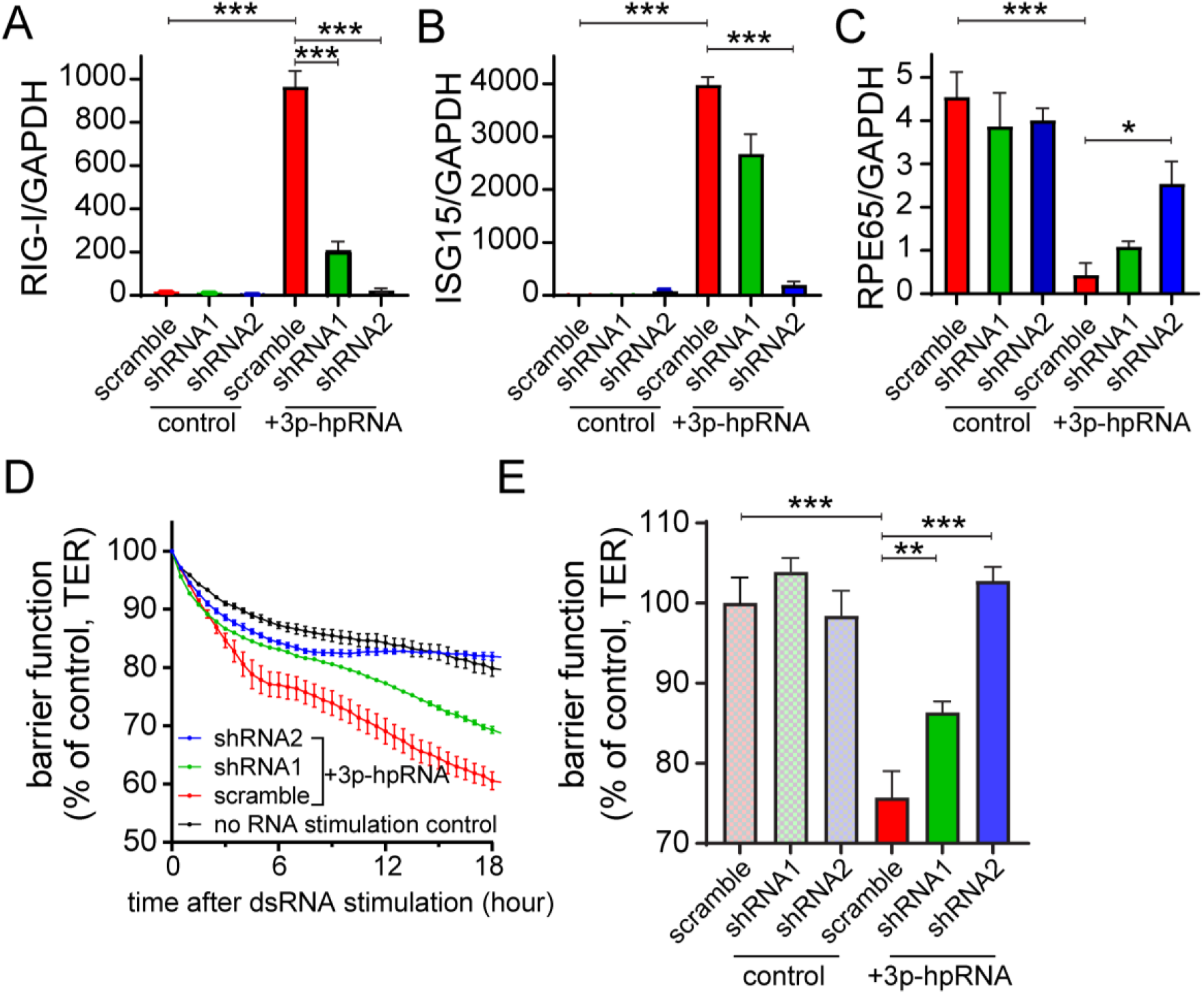
Synthetic dsRNA induces RIG-I dependent interferon response in RPE. (A-B) RT-qPCR analysis of iPS-RPE transcripts *RIG-I, ISG15*, and *RPE65* after shRNA knockdown of *RIG-I* with or without exposure to 3p-hpRNA. \**p*<0.05 and \*\*\**p*<0.001, Student’s t-test (n=3). (D) Representative barrier function analysis measured through TER, relative to timepoint zero, of iPS-RPE exposed to 3p-hpRNA with or without shRNA knockdown of *RIG-I* (n=3). (E) Cumulative analysis (N=3) at timepoint 18 h of barrier function measured through TER, relative to timepoint zero, of iPS-RPE exposed to 3p-hpRNA with or without shRNA knockdown of *RIG-I*. \*\**p*<0.01 and \*\*\**p*<0.001, Student’s t-test.

H. Kaneko *et al*. previously showed that the retrotransposon *Alu*, a noncoding dsRNA, is abundant in RPE during GA [8, 16]. *Alu* accumulation has been shown to induce RPE degeneration both *in vitro* and *in vivo* in a RIG-I independent manner [8, 17]. To evaluate if *Alu* dsRNA (*dsAlu*) could also induce RIG-I activation, we generated, purified, and annealed *in vitro* transcribed *dsAlu* and transfected into iPS-RPE with or without shRNA knockdown of RIG-I. For this study, we utilized only the RIG-I targeting shRNA2 for its superior efficiency (Figure 1A). We found that *dsAlu* was able to significantly induce both RIG-I and ISG15 expression and repress RPE65 expression in non-targeting controls but not in RIG-I-silenced cells (Figure 2A-C). Additionally, *dsAlu* transfection also induced significant TER loss, which was recovered with RIG-I knockdown (Figure 2D-E).

**Figure 2:**
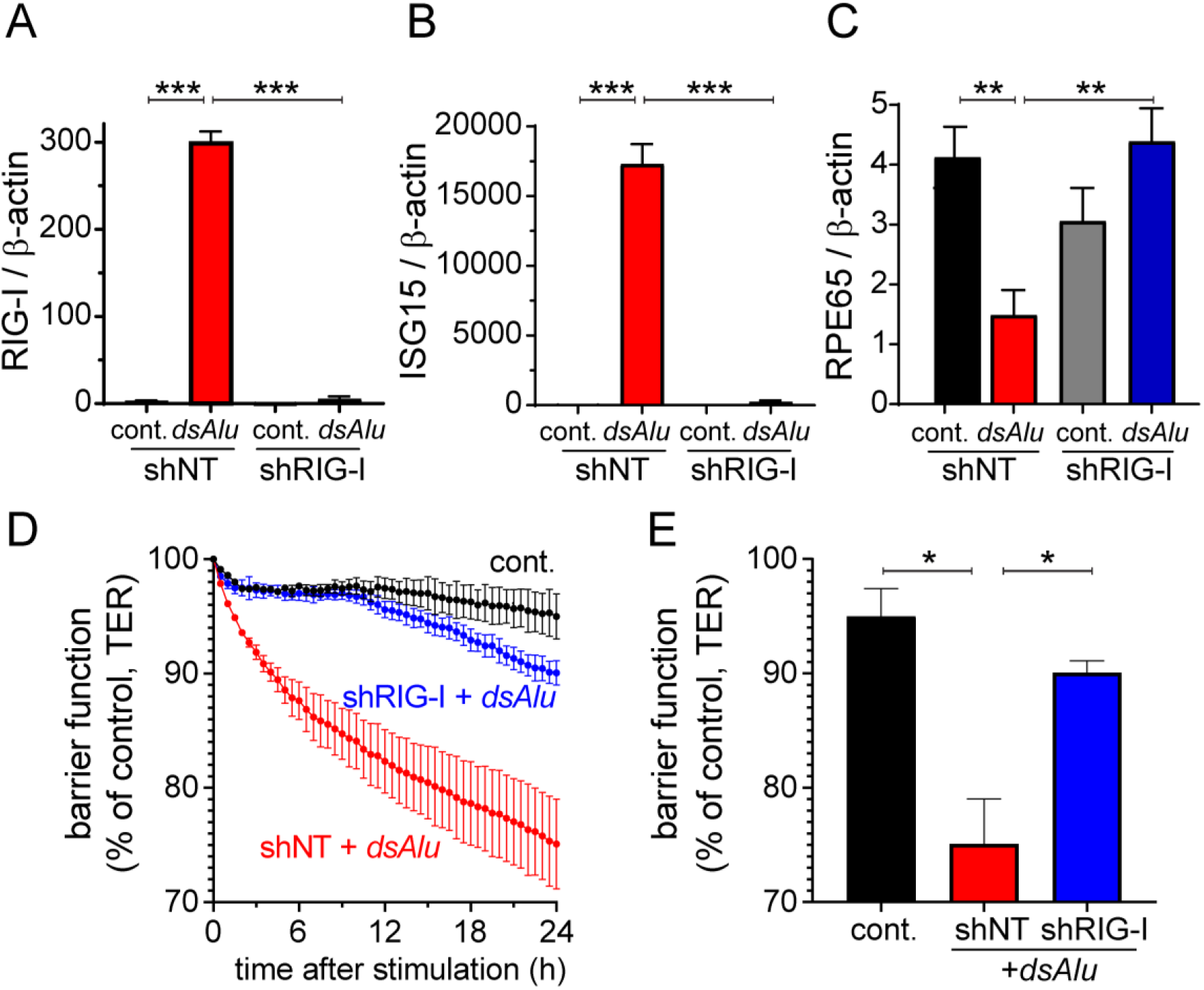
Double-stranded *Alu* induces RIG-I interferon response in RPE. (A-C) RT-qPCR analysis of iPS-RPE transcripts *RIG-I* (A), *ISG15* (B), and *RPE65* (C) after shRNA knockdown of *RIG-I* with or without exposure to *dsAlu*. \*\**p*<0.01 and \*\*\**p*<0.001, Student’s t-test (n=4). (D) Representative barrier function analysis measured through TER, relative to timepoint zero, of iPS-RPE exposed to *dsAlu* with or without shRNA knockdown of *RIG-I* (n=4). (E) Cumulative analysis of (N=3) barrier function at timepoint 24 h measured through TER, relative to timepoint zero, of iPS-RPE exposed to *dsAlu* with or without shRNA knockdown of *RIG-I*. \**p*<0.05, Student’s t-test.

### dsRNA accumulates in AMD donor RPE

We have previously shown that RIG-I is upregulated in the RPE with increasing progression of AMD [6]. Our findings using 3p-hpRNA and *dsAlu* suggest that dsRNA accumulation and RIG-I signaling may be relevant contributors to AMD-associated inflammation [18]. To further support a pathogenic role for dsRNA in inflammation associated with AMD progression, we stained donor ocular samples using a dsRNA specific antibody (J2, Scions) [19]. Post-mortem eyes from 4 aged-matched non-AMD donors, 7 intermediated AMD (iAMD) donors, and 7 GA donors were processed and cross-sectioned for immunohistochemistry (IHC) (Figure 3A-C). Macular dsRNA staining was compared and scored between AMD stages. Our results revealed a striking accumulation of dsRNA in the RPE that increased with AMD progression (Figure 3). Putting together the increased RIG-I protein levels, type I interferon expression, and dsRNA accumulation during AMD progression, we propose that dsRNA accumulation leads to cellular degeneration through a RIG-I mediated type I interferon response.

**Figure 3:**
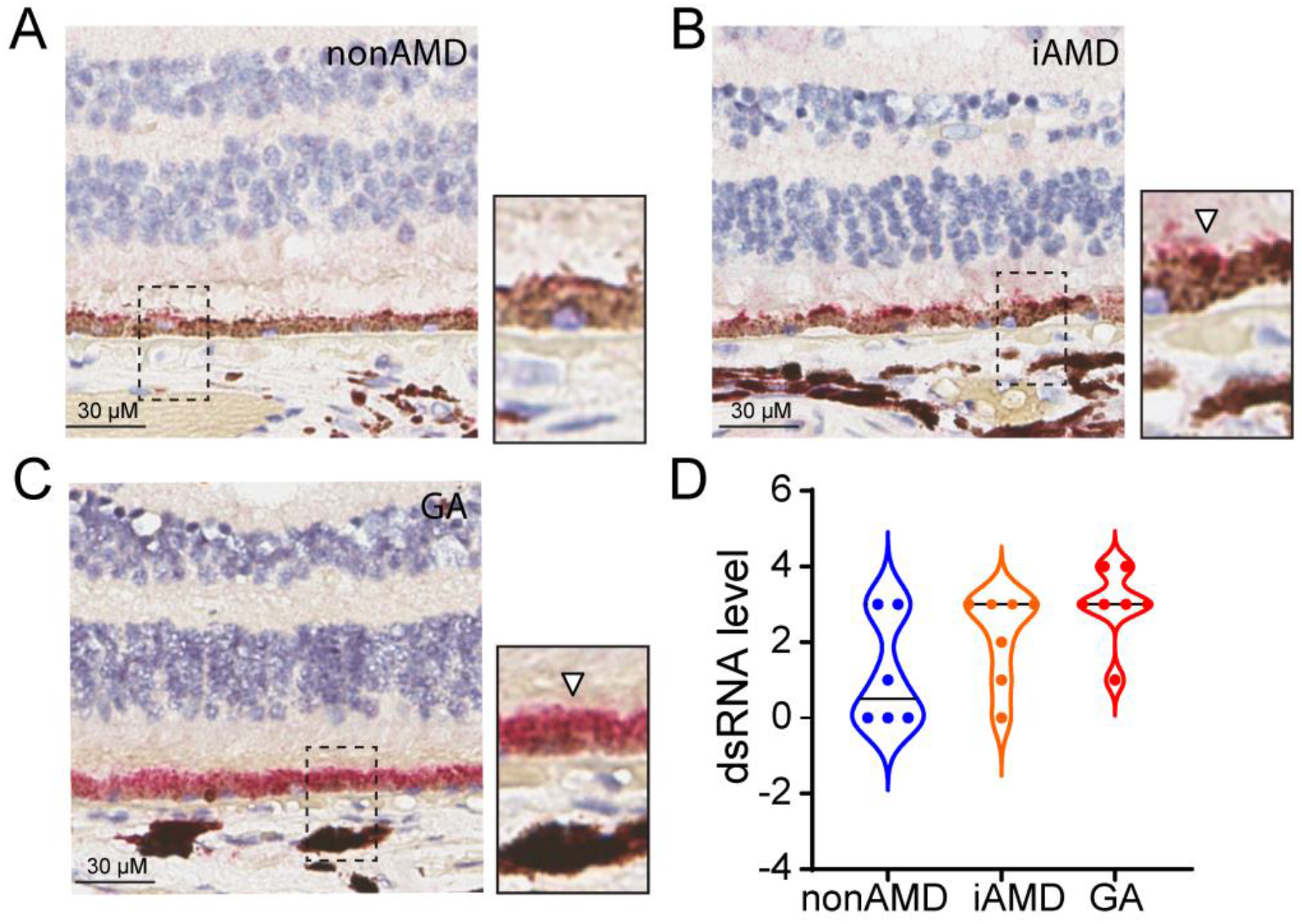
RPE accumulate dsRNA with AMD progression. (A-C) Representative immunohistochemistry for dsRNA of human donor eyes with intermediate AMD (iAMD, B), geographic atrophy (GA, C), or age-matched controls (non-AMD, A). Red = dsRNA. Blue = Hematoxylin. Brown = pigment. Dotted line indicates zoomed-in region pop-out. White arrowheads indicate positive staining in RPE. (D) Qualitative analysis for dsRNA staining intensities. Each dot represents individual scored patient sample within indicated stage (nonAMD N=6, iAMD N=7, GA N=7).

### Profiling of 3p-hpRNA exposed iPS-RPE identifies response pathways

The synthetic dsRNA-mimetic 3p-hpRNA induces the strongest RIG-I mediated type I interferon response, as well a degenerative phenotype in iPS-RPE cells *in vitro*. To confirm our mechanism of dsRNA-dependent induction of RPE degeneration *in vivo*, we designed an acute model for dsRNA accumulation in the back of the eye through subretinal injection of 3p-hpRNA.

Previous reports have utilized NeuroPorter (Millipore) as a method for efficient delivery of nucleic acids to the RPE through subretinal injection [20]. We first validated the efficiency of NeuroPorter as a transfection agent for dsRNA using fluorescently conjugated 3p-hpRNA on iPS-RPE (Figure 4A). We then generated a dose-response of 3p-hpRNA on *ISG15* expression to define the maximal effective dose (MED), which was 40 ng/ml (Figure 4B). To better understand the effects of RIG-I activation on RPE health, we evaluated potential changes in markers of RPE identity and epithelial health following 3p-hpRNA transfection. Using a Gene Array, we analyzed changes in RPE signature and epithelial-to-mesenchymal transition (EMT) genes in response to increasing doses of 3p-hpRNA up to the MED (Figure 4C) [21] and observed a reduction of multiple RPE specific genes (*RPE65, LRAT, BEST1, RDH5*) with 3p-hpRNA treatment, while EMT transcripts (*SERPINE1, ZEB1, TGFB1, VIM*) increased. Reduction of RPE65 suggested dedifferentiation and EMT which has been previously associated with AMD [22].

**Figure 4:**
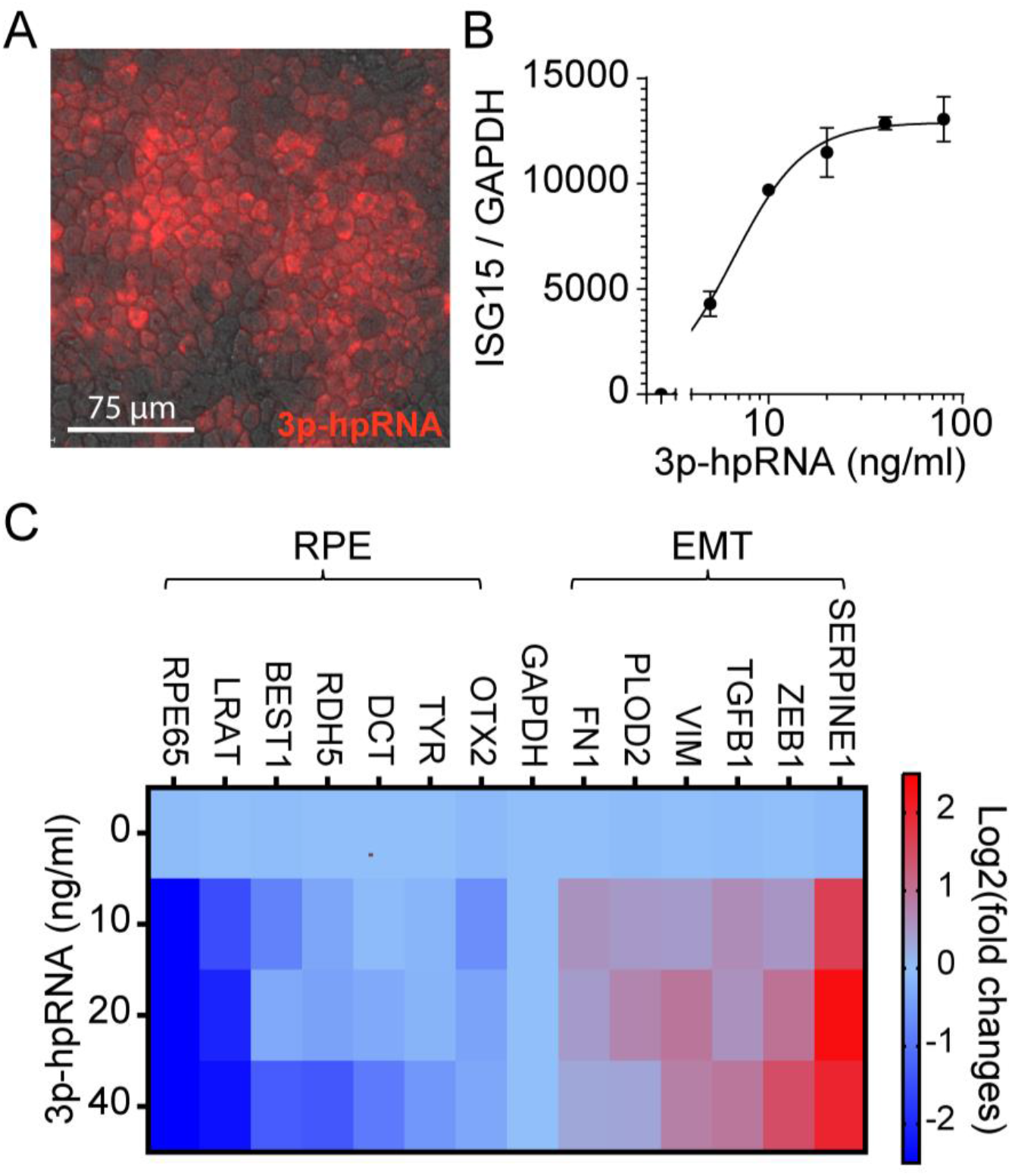
*In vitro* pharmacokinetic and pharmacodynamics for 3p-hpRNA transfection. (A) iPS-RPE transfected with A647-3p-hpRNA using Neuroporter. Red = AF647-3p-hpRNA (n=2). (B) Pharmacokinetic response of iPS-RPE to 3p-hpRNA exposure, measured through *ISG15* expression (n=4). (C) Gene array analysis (n=4) of 3p-hpRNA exposed iPS-RPE for RPE marker genes and EMT marker genes. Heat map indicates upregulation of expression relative to vehicle treated (0 ng/ml group), red = upregulated, light blue = no change, dark blue = downregulated.

### Development of acute *in vivo* RIG-I activation model

Following the optimization of 3p-hpRNA delivery *in vitro* we applied a similar approach *in vivo* to better assess the biological consequences of dsRNA accumulation to RPE. AF647-3p-hpRNA (2 μg/eye) or vehicle were delivered by subretinal injection to adult C57Bl/6J mice and evaluated 24 h post injections. Effective uptake of 3p-hpRNA in RPE was ascertained by immunolabelling of RPE flatmounts with the actin filament stain, phalloidin. Imaging 200 μm from the injection site showed accumulation of AF647-3p-hpRNA in RPE, confirming widespread *in vivo* transfection (Figure 5A).

**Figure 5:**
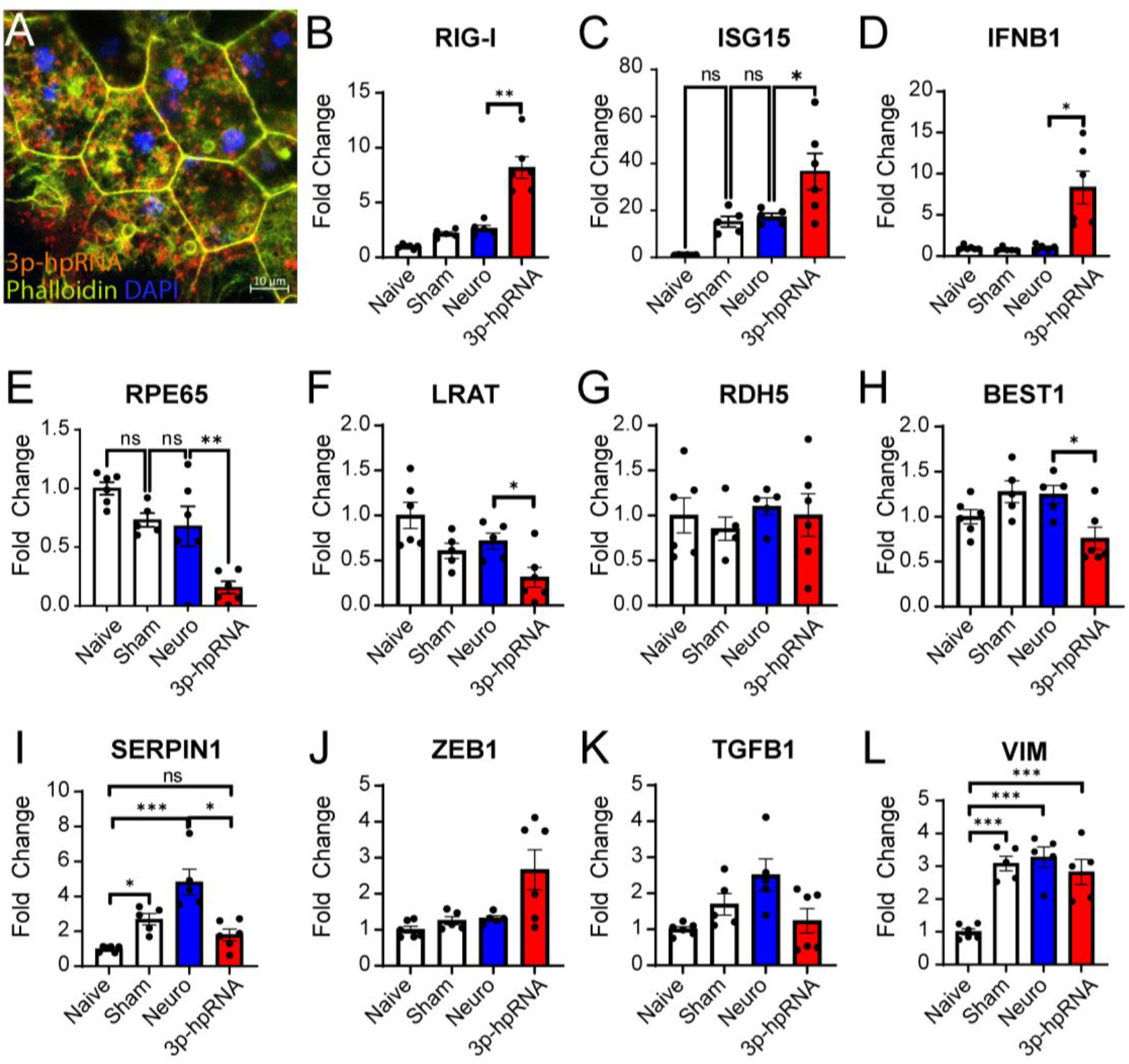
*In vivo* pharmacokinetic and pharmacodynamics of 3p-hpRNA subretinal injection. (A) Representative flatmount imaging of AF647-3p-hpRNA 24 h after subretinal injection, co-stained with ZO-1 to identify RPE. Red = 3p-hpRNA, Green = phalloidin, Blue = DAPI. Scale bar = 10 μm. (B-L) RT-qPCR analysis of isolated PEC 24 h after subretinal injection of 3p-hpRNA for RIG-I response genes (B-D), RPE function and identity genes (E-H), and EMT marker genes (I-L). \**p*<0.05, \*\**p*<0.01, \*\*\**p*<0.001 and ns, not significant, ANOVA.

We next assessed the effect of subretinal injection of 3p-hpRNA on the type I interferon response and RPE transcripts. Within 24 h post-injection, substantial induction of *ISG15, IFNβ1*, and *RIG-I* was measured, indicating activation of RIG-I (Figure 5B-D). Analysis of target genes identified by the Gene Array revealed a significant decrease in key RPE characteristic genes (*RPE65, LRAT, BEST1*) (Figure 4D-G). However, contrary to our *in vitro* data, no changes in EMT marker genes (*SERPIN1, ZEB1, TGFB1, VIM*) (Figure 4H-K) were detected post-3p-hpRNA transfection. Additional timepoints were analyzed (7 and 14-days after 3p-hpRNA injection) (Supplemental Figure 1) and showed that *RIG-I, IFNβ1* and *RPE65* returned to baseline levels by day 7 post-injection. This indicates that the decrease of RPE markers and increase in interferon response genes was acute and could return to baseline levels over time.

### dsRNA induces RPE degeneration & inflammation *in vivo*

Although significant changes in relevant signature genes were measured after delivery of 3p-hpRNA to RPE *in vitro* and *in vivo*, no marked morphological anomalies indicative of cell stress or tissue degeneration were observed by 24 h after ZO-1 staining (data not shown). We repeated the injection with a higher dose of 3p-hpRNA (6 μg/eye) and performed morphological assessment of RPE health through immunostaining of ZO-1. Using a grading system, participants could observe significant changes in RPE morphology, including both cellular swelling, shrinking, and rounding (Supplemental Figure 2). We additionally confirmed that changes to signature genes were comparable to the low dose injection at 72 h (Supplemental Figure 3). The grading consensus from 3 independent graders indicates that the 3p-hpRNA injection leads to clear pathological effects on RPE morphology, characterized by an increase in hyper- and hypo-morphological features (Figure 6C). The grading results were confirmed by automatized quantification of cell roundness, a widely used RPE morphometric parameter [23], indicating abnormalities (Figure 6D).

**Figure 6:**
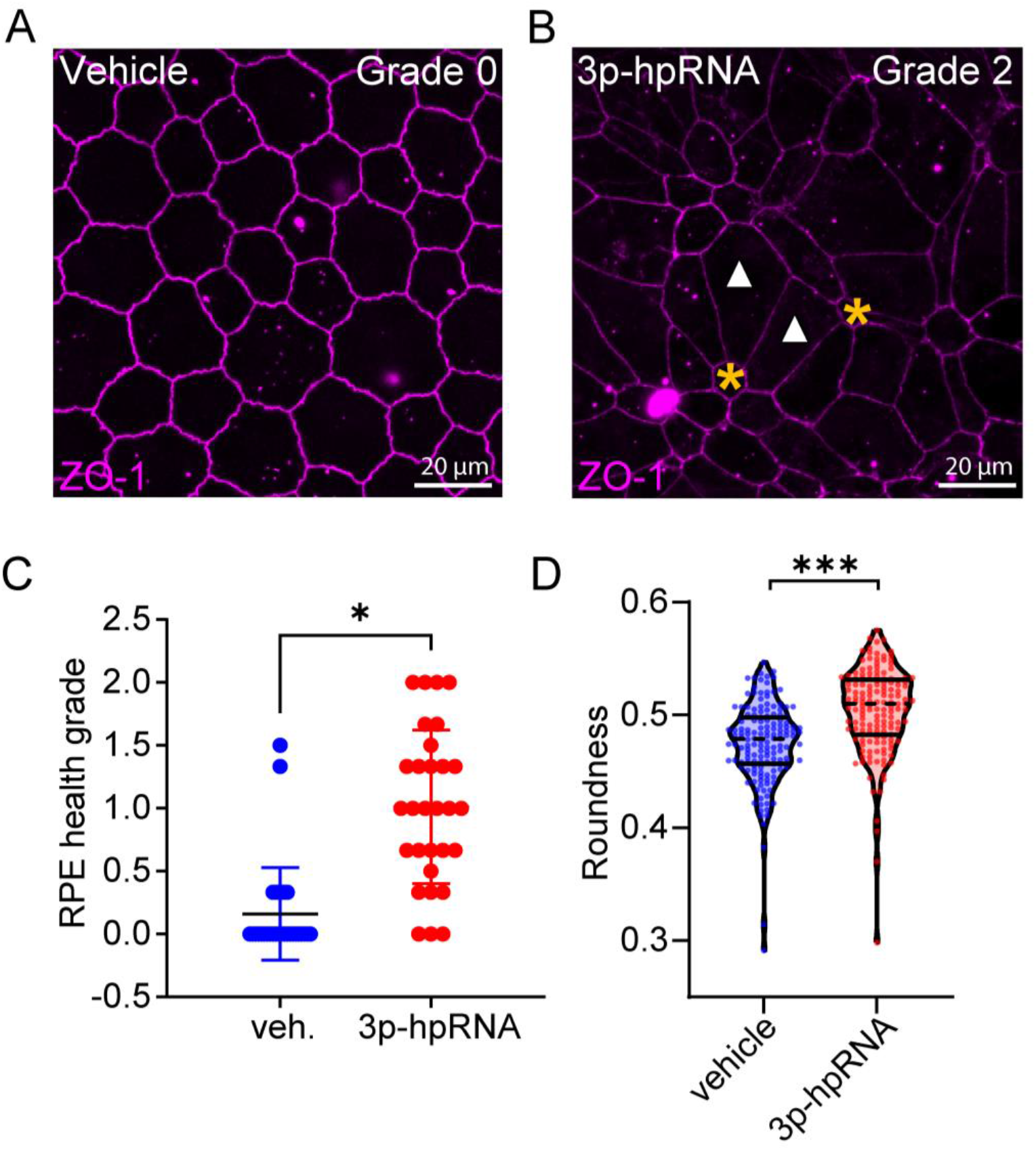
3p-hpRNA induces acute RPE stress *in vivo*. (A-B) ZO-1 immunostaining of flatmounts 72 h after subretinal injection of vehicle or 3p-hpRNA. White arrowheads indicate cellular swelling or enlargment. Yellow asterisks indicate cellular skrinkage. Purple = ZO-1. (C) Averaged qualitative analysis of RPE health using ZO-1 immunostaining from three independent graders. \**p*<0.05, Welch’s t-test. (D) Qualitative analysis of cell roundness from ZO-1 immunostaining. \*\*\**p*<0.001, Student’s t-test.

Morphological changes were further characterized using transmission electron microscopy (TEM). Using vehicle injected animals as controls, observations were conducted approximately 300 to 500 μm away from the subretinal injection site as to avoid areas affected by any procedure-related effects. In 3p-hpRNA-injected animals, ultrastructural evidence of photoreceptor outer segment (POS) degeneration and necrosis, RPE alterations (hypertrophy, increased phagosomes) and inflammatory cell infiltration were observed (Figure 7A-B), validating that RIG-I activation was detrimental for retinal health. The observation of inflammatory cell infiltration was particularly interesting, in conjunction with POS degeneration, due to our previous reports detailing the degenerative influence of macrophage activation on retinal health [14]. Because secretion of IFNβ can cause recruitment and activation of phagocytes such as macrophages [24], we measured macrophage activation though *IBA1* expression after 3p-hpRNA transfection and found a significant increase in expression (Figure 7C). Immunostaining of RPE flatmounts confirmed macrophage recruitment and activation to regions surrounding 200-300 μm from the injection site with 3p-hpRNA, but not in vehicle (Figure 7D-E). Increased detection of IBA1+ cells was also observed 72 h post-injection (Figure 7F).

**Figure 7:**
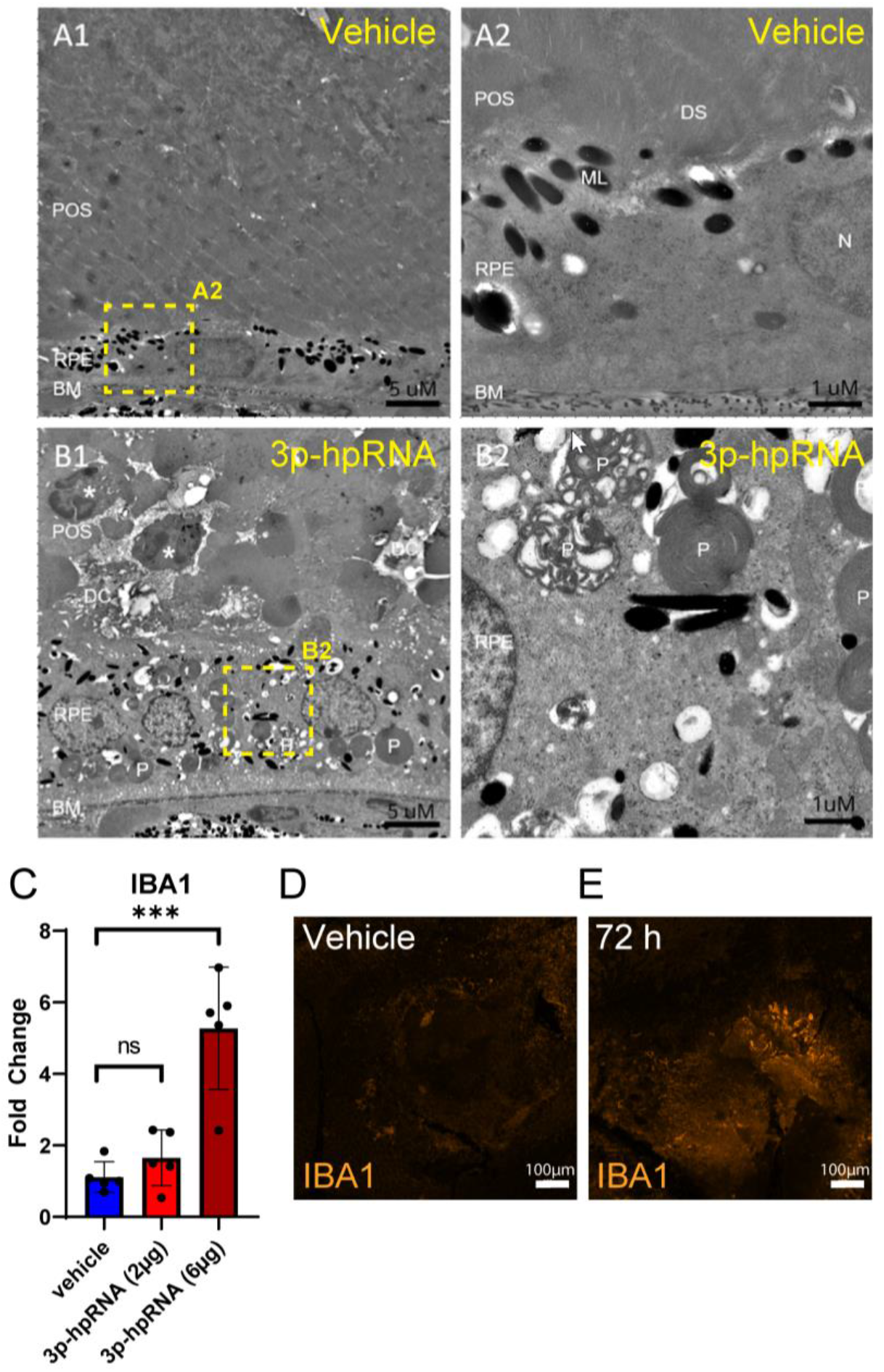
3p-hpRNA subretinal injection to RPE induces lymphocyte recruitment and activation. Transmission electron micrographs of the deep layers of the retina 300-500 μm distant from the edge of the subretinal injection site in 72 h after animals injected with control vehicle transfection agent (A1 low magnification, A2 higher magnification) or 3p-hpRNA dsRNA (B1 low magnification, B2 higher magnification). Yellow dotted lined indicates high magnification region within low magnification image. POS: photoreceptor outer segments; RPE: retinal pigmented epithelium; BM: Bruch’s membrane; P: phagosomes; Star: inflammatory cells; ML: melanosome/lipofuscin; N: nucleus; DS: disk stacks; DC: degenerated and necrotic cells. (C) RT-qPCR analysis for *IBA1* of isolated PEC from subretinal injection of vehicle or low- and high-doses of 3p-hpRNA. \*\*\**p*<0.001, and ns, not significant, ANOVA (D-E) IBA1 immunostaining of PEC flatmounts post subretinal injection of vehicle (D) or 3p-hpRNA (E) after 72 h. Orange = IBA1.

## Discussion

Over the past few years, a better understanding of the role and origin of endogenous dsRNA in human disease has had great appreciation. Activation of RLR (RIG-I-like receptor) via dsRNA can promote the secretion of inflammatory cytokines such as IFN, subsequently recruiting leucocytes [7]. While useful for combating acute infections, hyperactivation of RLR through endogenously derived dsRNA can lead to chronic inflammation. During AMD, RPE cells show a strong signature for IFN, leading us to investigate the mechanisms which this signature response arises from. Previously, we have shown that activation RIG-I is the predominant driver of IFN in RPE cells [6]. Here, we complement our findings with a mechanistic investigation of the molecular drivers of RIG-I activation in RPE and identify dsRNA accumulation as a likely initiator of IFN signaling in AMD.

Many potential mechanisms to induce dsRNA accumulation have been described, including misregulation of RNA processing enzymes like DICER1 and ADAR1 [8, 25]. IHC for dsRNA in AMD patient samples indicated that dsRNA accumulated in RPE as early as iAMD, and that this accumulation continued to increase until the GA stage. The reports from H. Kaneko *et al*. showed that the retrotransposable element *Alu* RNA was enriched in GA [8]. *Alu* RNA has been shown to elicit an NLRP3-dependent inflammasome response, as well as RPE degeneration though L1 reverse transcription into *Alu* cDNA and activation of cGAS [26, 27]. In contrast to these findings, we have found that cGAS was not present in healthy RPE but rather limited to pro-inflammatory macrophages, failing to explain the progressive degenerative decline of RPE health through the early stages of AMD [6]. The data presented here indicates that the deleterious effects of *Alu* RNA can be phenocopied through synthetic dsRNA-mimetic 3p-hpRNA; and regardless of the identity of the dsRNA involved, loss of the RNA sensor RIG-I led to recovery. Many questions remain on the upstream mechanisms for dsRNA accumulation, but these results spark a revisit of the proposed role of *Alu* RNA in AMD progression.

While we have focused on the RPE as the primary culprit for retinal degeneration in this study, the photoreceptors may also be a major contributor. A key function of the RPE involves clearing the photoreceptor outer segments (OS). The OS can become highly oxidized, and contain toxic metabolites that can lead to the activation of pro-inflammatory pathways in the RPE [28]. There may be potential dsRNA inducers found within damaged or oxidized OS, or failure to clear these volatile segments may lead to damage associated with dsRNA accumulation [29].

The involvement of dsRNA has been reported in other diseases such as cancer [30], Huntington’s Disease [31], osteoarthritis (OA) [32], alcohol-associated liver disease (ALD) [33], and chronic kidney disease (CKD) [34], to name a few. In the case of ALD, OA, and CKD, mitochondrial dysfunction has been implicated in the release of mitochondrial dsRNA to activate a PKR mediated IFN response. Mitochondrial dysfunction is characteristic of AMD [35], and it has been shown that dsRNA released from mitochondria may initiate RLR signaling [36]. Alternatively, nuclear transcription of dsRNA due to epigenetic changes, such as in the case of cancer and Huntington’s disease, can produce immunogenic LINEs and SINEs, such as *Alu*. The identity and source of dsRNA found in RPE during AMD still needs exploration.

Regardless, exposure of RPE cells to dsRNA *in vitro* led to an increase in IFN related genes. Several indicators pointed to an EMT-like phenotype after exposure, including loss of barrier function and an increased expression in EMT marker genes such as *SERPINE1* [21]. Interestingly though, dsRNA challenge in our *in vivo* model did not trigger a similar EMT signature. Several possibilities exist as to why this might have occurred. The transcriptomic analyses of our preclinical samples were performed using RNA isolated from the entire PEC, thus not restricted to RPE cells, nor to the region with the highest dsRNA transfection, which may have diluted the RPE signal below detection. Another possibility is that a molecular EMT signature can only be observed under chronic stress conditions. Findings by Zhao *et al*. show that while loss of RPE identity markers such as RPE65 can occur early after metabolic dysregulation induced stress *in vivo*, induction of canonical EMT marker genes can take several weeks to be detectable [37]. Interestingly, they did observe early hypertrophic morphology changes, similar to the ones observed in our *in vivo* model of retinal dsRNA challenge, preceding induction of an EMT signature. This suggests that an EMT signature may still be inducible by dsRNA, but may require sustained dsRNA delivery to accumulate in RPE. Furthermore, the animals used in this study were young and healthy. It is not known whether the capacity of RPE to clear dsRNA is affected by age or disease state, so repeating these studies on aged or diseased animal models would be of interest in further establishing a mechanistic link between dsRNA accumulation, RIG-I, and EMT in RPE.

Most importantly, the *in vivo* model developed here exemplifies the importance of dsRNA regulation in ocular health. Delivery of dsRNA led to a potent IFN response and degeneration of both RPE and POS, as observed both with immunostaining and TEM. Because the POS and the RPE are functionally interdependent [38], it is difficult to postulate from the TEM observations at a single timepoint whether the RPE change are secondary to the photoreceptor degeneration or from a different source entirely, such as inflammation. While these primary RIG-I mediated effects are acute, several indicators point to lingering inflammatory phenotype, such as recruitment and activation of macrophages. Previous work from our group has found that macrophages could contribute to RPE and photoreceptor degeneration [14]. Clearance of dsRNA may be a rapidly occurring process, and as indicated by the work exploring DICER1, may protect cells from long-term detrimental effects [16]. To this end, chronic exposure of dsRNA may lead to an even more dramatic response. Further expansion of this model, such as through knocking-down DICER1, may further progress the phenotype. Additionally, creating a chronic model through spontaneous endogenous accumulation of dsRNA in RPE would answer questions regarding the long-term consequence of chronic RIG-I activation.

Overall, we conclude that RIG-I mediated signaling through detection of dsRNA may be a key pathway driving AMD progression. We showed the essential role of RIG-I in responding to dsRNA *in vitro*, and IHC analysis of donor samples found that dsRNA was highly prevalent in the RPE during iAMD and GA. Finally, acute exposure of RPE to dsRNA via transfection of 3p-hpRNA subretinally led to RIG-I activation, RPE stress, and inflammation characteristic of AMD. This signaling model could be useful for better understanding AMD progression, and uncover new avenues for intervention.

## Acknowledgements

We would like to thank Omar Delgado for assisting with euthanasia during *in vivo* studies. We would also like to thank Megan Serpa for validating the dsRNA antibody J2.

## Author Contributions

The authors confirm contributions to the manuscript as follows: conceptualization: Y.B., J.A. methodology: G.K., Y.X., K.B., J.S., J.M., M.T., H.H., M.K.P., M.C., J.D. validation: G.K., Y.X., K.B., H.H. formal analysis: G.K., Y.X., K.B. investigation: G.K., Y.X., K.B., X.W., J.M., C.H. resources: J.R.G., Y.Q., R.E., Q.H., G.P., M.S.G. data curation: Y.B., J.A. writing – original draft: K.B., Y.B. writing- review & editing: K.B., G.K. Y.X., J.R.G., M.S.G, J.A., Y.B. visualization: G.K., Y.X., K.B., J.R.G., J.A., Y.B. supervision: Y.B., J.A., Q.H., G.P., M.S.G. project administration: Y.B., J.A., Q.H., G.P., M.S.G. All authors reviewed the results and approved the final version of the manuscript.

## References

1. Kauppinen, A., et al., Inflammation and its role in age-related macular degeneration. Cell Mol Life Sci, 2016. 73(9): p. 1765–86.

2. Boyer, D.S., et al., The Pathophysiology of Geographic Atrophy Secondary to Age-related Macular Degeneration and the Complement Pathway as a Therapeutic Target. Retina, 2017. 37(5): p. 819–835.

3. Liao, D.S., et al., Complement C3 Inhibitor Pegcetacoplan for Geographic Atrophy Secondary to Age-Related Macular Degeneration: A Randomized Phase 2 Trial. Ophthalmology, 2020. 127(2): p. 186–195.

4. Wei, T.T., et al., Interferon-γ induces retinal pigment epithelial cell Ferroptosis by a JAK1-2/STAT1/SLC7A11 signaling pathway in Age-related Macular Degeneration. Febs j, 2022. 289(7): p. 1968–1983.

5. David R. Guyer, M.A.P.A., MD; Lawrence, et al., Interferon alfa-2a is ineffective for patients with choroidal neovascularization secondary to age-related macular degeneration. Results of a prospective randomized placebo-controlled clinical trial. Pharmacological Therapy for Macular Degeneration Study Group. Arch Ophthalmol, 1997. 115(7): p. 865–72.

6. Schustak, J., et al., Mechanism of Nucleic Acid Sensing in Retinal Pigment Epithelium (RPE): RIG-I Mediates Type I Interferon Response in Human RPE. Journal of Immunology Research, 2021. 2021: p. 9975628.

7. De Cecco, M., et al., L1 drives IFN in senescent cells and promotes age-associated inflammation. Nature, 2019. 566(7742): p. 73–78.

8. Kaneko, H., et al., DICER1 deficit induces Alu RNA toxicity in age-related macular degeneration. Nature, 2011. 471(7338): p. 325–330.

9. Häsler, J. and K. Strub, Alu elements as regulators of gene expression. Nucleic Acids Res, 2006. 34(19): p. 5491–7.

10. Aune, T.M., et al., Alu RNA Structural Features Modulate Immune Cell Activation and A-to-I Editing of Alu RNAs Is Diminished in Human Inflammatory Bowel Disease. Front Immunol, 2022. 13: p. 818023.

11. Sarks, S.H., Ageing and degeneration in the macular region: a clinico-pathological study. Br J Ophthalmol, 1976. 60(5): p. 324–41.

12. Olsen, T.W. and X. Feng, The Minnesota Grading System of eye bank eyes for age-related macular degeneration. Invest Ophthalmol Vis Sci, 2004. 45(12): p. 4484–90.

13. Ferris, F.L., 3rd, et al., Clinical classification of age-related macular degeneration. Ophthalmology, 2013. 120(4): p. 844–51.

14. Twarog, M., et al., TNFα induced by DNA-sensing in macrophage compromises retinal pigment epithelial (RPE) barrier function. Sci Rep, 2023. 13(1): p. 14451.

15. Hochheiser, K., et al., Cutting Edge: The RIG-I Ligand 3pRNA Potently Improves CTL Cross-Priming and Facilitates Antiviral Vaccination. J Immunol, 2016. 196(6): p. 2439–43.

16. Kim, Y., et al., DICER1/Alu RNA dysmetabolism induces Caspase-8-mediated cell death in age-related macular degeneration. Proc Natl Acad Sci U S A, 2014. 111(45): p. 16082–7.

17. Kleinman, M.E., et al., Short-interfering RNAs induce retinal degeneration via TLR3 and IRF3. Mol Ther, 2012. 20(1): p. 101–8.

18. Datta, S., et al., The impact of oxidative stress and inflammation on RPE degeneration in non-neovascular AMD. Prog Retin Eye Res, 2017. 60: p. 201–218.

19. Schönborn, J., et al., Monoclonal antibodies to double-stranded RNA as probes of RNA structure in crude nucleic acid extracts. Nucleic Acids Res, 1991. 19(11): p. 2993–3000.

20. Kachi, S., et al., Nonviral ocular gene transfer. Gene Ther, 2005. 12(10): p. 843–51.

21. Sripathi, S.R., et al., Proteome Landscape of Epithelial-to-Mesenchymal Transition (EMT) of Retinal Pigment Epithelium Shares Commonalities With Malignancy-Associated EMT. Mol Cell Proteomics, 2021. 20: p. 100131.

22. Cao, D., et al., Hyperreflective Foci, Optical Coherence Tomography Progression Indicators in Age-Related Macular Degeneration, Include Transdifferentiated Retinal Pigment Epithelium. Invest Ophthalmol Vis Sci, 2021. 62(10): p. 34.

23. Bhatia, S.K., et al., Analysis of RPE morphometry in human eyes. Mol Vis, 2016. 22: p. 898–916.

24. Kopitar-Jerala, N., The Role of Interferons in Inflammation and Inflammasome Activation. Frontiers in Immunology, 2017. 8.

25. Solomon, O., et al., RNA editing by ADAR1 leads to context-dependent transcriptome-wide changes in RNA secondary structure. Nature Communications, 2017. 8(1): p. 1440.

26. Tarallo, V., et al., DICER1 loss and Alu RNA induce age-related macular degeneration via the NLRP3 inflammasome and MyD88. Cell, 2012. 149(4): p. 847–59.

27. Fukuda, S., et al., Alu complementary DNA is enriched in atrophic macular degeneration and triggers retinal pigmented epithelium toxicity via cytosolic innate immunity. Sci Adv, 2021. 7(40): p. eabj3658.

28. Hoppe, G., et al., Oxidized low density lipoprotein-induced inhibition of processing of photoreceptor outer segments by RPE. Invest Ophthalmol Vis Sci, 2001. 42(11): p. 2714–20.

29. Panfoli, I., et al., Proteomic analysis of the retinal rod outer segment disks. J Proteome Res, 2008. 7(7): p. 2654–69.

30. Kang, M., et al., Double-stranded RNA induction asa potential dynamic biomarkerfor DNA-demethylating agents. Mol Ther Nucleic Acids, 2022. 29: p. 370–383.

31. Peel, A.L., et al., Double-stranded RNA-dependent protein kinase, PKR, binds preferentially to Huntington’s disease (HD) transcripts and is activated in HD tissue. Hum Mol Genet, 2001. 10(15): p. 1531–8.

32. Kim, S., et al., Mitochondrial double-stranded RNAs govern the stress response in chondrocytes to promote osteoarthritis development. Cell Rep, 2022. 40(6): p. 111178.

33. Lee, J.H., et al., Mitochondrial Double-Stranded RNA in Exosome Promotes Interleukin-17 Production Through Toll-Like Receptor 3 in Alcohol-associated Liver Injury. Hepatology, 2020. 72(2): p. 609–625.

34. Zhu, Y., et al., Polynucleotide phosphorylase protects against renal tubular injury via blocking mt-dsRNA-PKR-eIF2α axis. Nat Commun, 2023. 14(1): p. 1223.

35. Tong, Y., Z. Zhang, and S. Wang, Role of Mitochondria in Retinal Pigment Epithelial Aging and Degeneration. Front Aging, 2022. 3: p. 926627.

36. Wiatrek, D.M., et al., Activation of innate immunity by mitochondrial dsRNA in mouse cells lacking p53 protein. Rna, 2019. 25(6): p. 713–726.

37. Zhao, C., et al., mTOR-mediated dedifferentiation of the retinal pigment epithelium initiates photoreceptor degeneration in mice. J Clin Invest, 2011. 121(1): p. 369–83.

38. Mazzoni, F., H. Safa, and S.C. Finnemann, Understanding photoreceptor outer segment phagocytosis: use and utility of RPE cells in culture. Exp Eye Res, 2014. 126: p. 51–60.

