## Supplemental Information for "Double-stranded RNA induces retinal pigment epithelium cell degeneration and inflammation"

| | Number | Average Age (years) |
|---|---|---|
| nonAMD | 6 (F=2, M=4) | 80.67 (F=78, M=86) |
| iAMD | 7 (F=4, M=3) | 85.43 (F=85, M=86) |
| GA | 7 (F=3, M=4) | 86.14 (F=88.25, M=83.33) |

**Supplemental Table 1: Patient sample summary used for IHC dsRNA analyses.** F = Female, M = Male, nonAMD = not diagnosed with AMD, iAMD = intermediate AMD, GA = Geographic Atrophy.

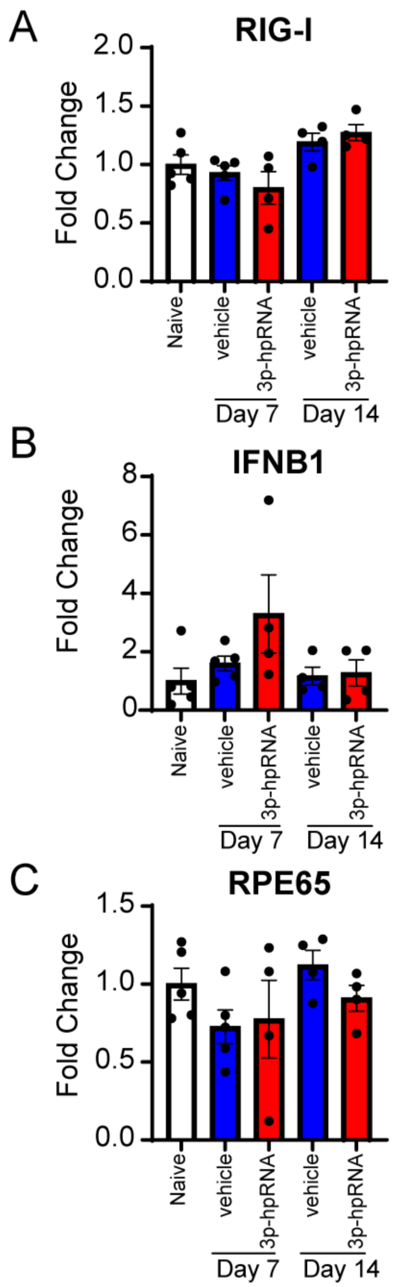


**Supplemental Figure 1: Extended time-course analysis after subretinal injection of low dose (2 µg) 3p-hpRNA.** (A-C) RT-qPCR analysis of *RIG-I* (A), *IFNβ1*(B), and *RPE65* (C) in posterior eye-cup (PEC) samples from 7 days and 14 days after 3p-hpRNA subretinal injection.

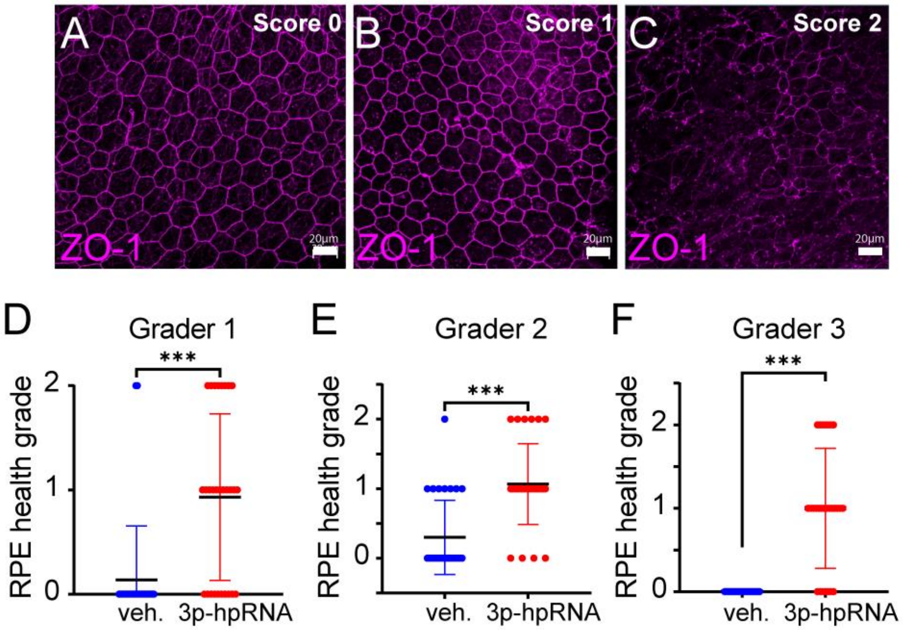


**Supplemental Figure 2: ZO-1 scoring of RPE flatmounts after subretinal injection of 3p-hpRNA (6 µg).** (A-C) Consensus representative images at different health scores among three individual graders. (D-F) Summary of scoring of individual graders ***$p$<0.001, Welch's t-test.

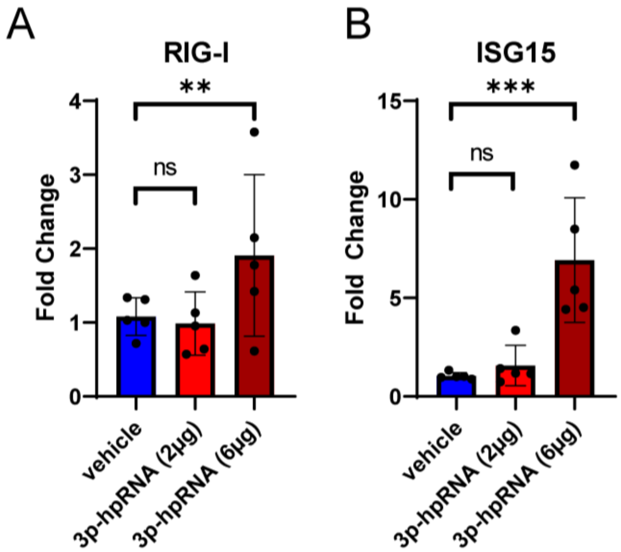


**Supplemental Figure 3: *In vivo* assay confirmation.** RT-qPCR analysis of vehicle, low-dose (2 µg), and high-dose (6 µg) of 3p-hpRNA at 72 h timepoint for *RIG-I* (A) and *ISG15* (B) in posterior eye-cup (PEC) samples. $^{**}p<0.01$, $^{***}p<0.001$, and ns, not significant, ANOVA.
